# Allochronic isolation between sympatric populations of an alpine butterfly

**DOI:** 10.64898/2025.12.01.691721

**Authors:** Christopher A. Halsch, Chris C. Nice, Katherine L. Bell, James A. Fordyce, Matthew L. Forister, Guillermo Garcia-Costoya, Arthur M. Shapiro, Eliza M. Grames

## Abstract

The role of isolation by distance in contributing to population structure across space is well understood, and the same principles should operate over time. Allochrony, the isolation of populations due to separation in time, can be a source of genetic differentiation, but remains infrequently documented between yearly asynchronous populations. In this study, we investigate divergence between two putative sympatric populations of the alpine butterfly *Oeneis chryxus ivallda* at Castle Peak, CA, USA. We find clear genetic differentiation between butterflies collected in odd and even years, with limited instances of admixture. The observed G_ST_ of 0.05 between the two populations is approximately equivalent to 26 km based on pairwise G_ST_ values observed between *O. chryxus* populations across space. These results demonstrate a potential source of population differentiation in systems that promote prolonged development, often found in high-elevation and high-latitude environments.

## Introduction

A central principle of evolutionary biology is that populations separated in space, subjected to varied selection pressures and stochastic processes, become differentiated (Wright 1943, Rousset 1997). Examples of ecological and evolutionary forces interacting to drive reproductive isolation across space are manifold (Slatkin 1987, Bohonak 1999, Rundle and Nosil 2005); however, these mechanisms can also operate over time (Hendry and Day 2005). Temporal, or allochronic, isolation can arise from phenological variance within a day, among seasons, or across years, but in each instance, populations separated in time become distinct (Fukami et al. 2003, Fudickar et al. 2016, Bell et al. 2017). The capacity of seasonal allochrony to contribute to divergence is relatively well established (Feder et al. 1988, Santos et al. 2007, Friesen et al. 2007), but the extent to which yearly temporal isolation, in which sympatric populations occur across years, can generate population structure is less understood (Taylor and Friesen 2017).

Unlike seasonal separation, where environments vary across time, yearly allochronic cohorts experience largely the same conditions, resulting in less variation for selection to act upon. Despite this, some instances of yearly allochrony promoting speciation are known, perhaps best exemplified by the highly periodic genus *Magicicada* (Ritchie 2001, Simon et al. 2022). Cicadas in this genus comprise three distinct species groups across eastern North America, each with species that emerge every 13 or 17 years and, as a result, are almost entirely reproductively isolated, co-occurring only every 221 years (Sota et al. 2013). Multiple hypotheses have been proposed to explain this phenomenon, ranging from temperature during the last glacial maximum (Ito et al. 2015) to predation pressure (Koenig and Liebhold 2013), both of which suggest that isolation results from selective pressure. Similar results are seen in Atlantic salmon (*Salmo salar*), where individuals return to their spawning grounds after spending between one and five years at sea (Hutchings and Jones 1998). The age at which fish return appears to be genetically determined, leading to reproductive isolation and divergence between sympatric populations with differing maturation ages (Johnston et al. 2014). In both systems, the influence of selection acting on asynchronous timing across years to establish and maintain reproductive isolation is clear.

While these cases demonstrate how selection can lead to substantial allochronic divergence, the capacity for stochastic processes to produce small-scale yearly temporal differentiation has been less explored, but may be common (Devaux and Lande 2008). Drift-induced allochronic isolation would be expected in environments that promote small, phenologically separated populations with short reproductive periods, as is often the case in insects (Heliövaara et al. 1994, Berlocher and Feder 2002), especially in cold, unpredictable environments characterized by low primary productivity or with low-quality food resources (Danks 1992). The pine bark bug (*Aradus cinnamomeus*) in southern Finland offers insight into this process, where two large populations are separated both spatially and by asynchronous two-year life cycles (Heliövaara and Väisänen 1987). Each large population has a much smaller sympatric population that emerges in the alternate year (Heliövaara and Väisänen 1987). These small, off-year populations are genetically distinct from the larger populations they co-occur with in space (Heliövaara et al. 1988). The two large populations, however, separated in space and time, are not genetically distinguishable from each other (Heliövaara et al. 1988), a pattern consistent with drift driving differentiation in the smaller populations. Similar questions have been studied in biennial alpine butterflies: *Erebia palarica* in Europe (Vila and Björklund 2004) and *Oeneis macounii* and *Oeneis melissa semidea* in North America (Gradish et al. 2015, 2019). In *O. melissa*, weak differentiation was detected between allochronic populations (Gradish et al. 2015), but in *E. palarica* and *O. macounii*, no difference was found. Thus, although prolonged development is widespread among insects in cold, high-latitude and high-elevation environments, the conditions under which it translates into measurable population structure remain unclear.

In this study, we investigate yearly allochronic separation between two putative sympatric populations of the alpine butterfly *Oeneis chryxus ivallda,* collected over five consecutive years from Castle Peak in the northern Sierra Nevada Mountains of California. Unlike many other biennial populations, which fluctuate between a high “on” year and low “off” year census population size, this locality has historically had a stable population size in both odd and even years, suggesting two distinct populations (Nice and Shapiro 2001, Halsch et al. 2024). Using a reduced-representation sequencing approach (Peterson et al. 2012), we explore whether there are differences between odd- and even-year butterflies collected at the same location. We then examine the admixture and evolutionary histories of both hypothesized populations. Finally, we quantify the magnitude of the observed differentiation by associating temporal with spatial differences. In doing so, we examine populations in an environment where genetic drift is likely the only factor contributing to divergence and explore a potential mechanism underlying differentiation in small, isolated, high-elevation insect populations.

## Methods

### Study system

*Oeneis chryxus* (Family: Nymphalidae, Subfamily: Satyrinae) is found in the northeastern United States, southeastern Canada, and above the tree line in montane western North America (Scott 1986). In California, *O. chryxus* has traditionally been considered two subspecies, *Oeneis chryxus ivallda* and *Oeneis chryxus stanislaus*; however, no genetic basis for this distinction has been identified (Hovanitz 1940, Porter and Shapiro 1989, MacDonald et al. 2024). Across the Sierra Nevada mountains, *O. chryxus* is primarily found in open, rocky alpine habitats, where it consumes grasses and sedges as larval host plants (Scott 1986). It develops over two years, and because of this, *O. chryxus* is abundant only in odd-numbered years in most locations within the Sierra (Nice and Shapiro 2001). However, in one locality, Castle Peak, they were found in high abundance every year.

### Data collection, extraction, and sequencing

Between 1989 and 1995, 527 *O. chryxus* individuals were collected from 13 sites across the Sierra Nevada and an additional site in the Great Basin as part of a separate study (Nice and Shapiro 2001) (Fig. 1). This collection effort included five consecutive years at Castle Peak, during which 17, 13, 25, 23, and 20 individuals were collected between 1991 and 1995, respectively. A smaller number of individuals were collected from the Sweetwater Mountains in 1992, 1993, and 1995 (50 total individuals, but only three from an even year). All individuals were originally processed for allozyme analysis, and homogenates were stored at -80°C after their use.

**Figure 1.**
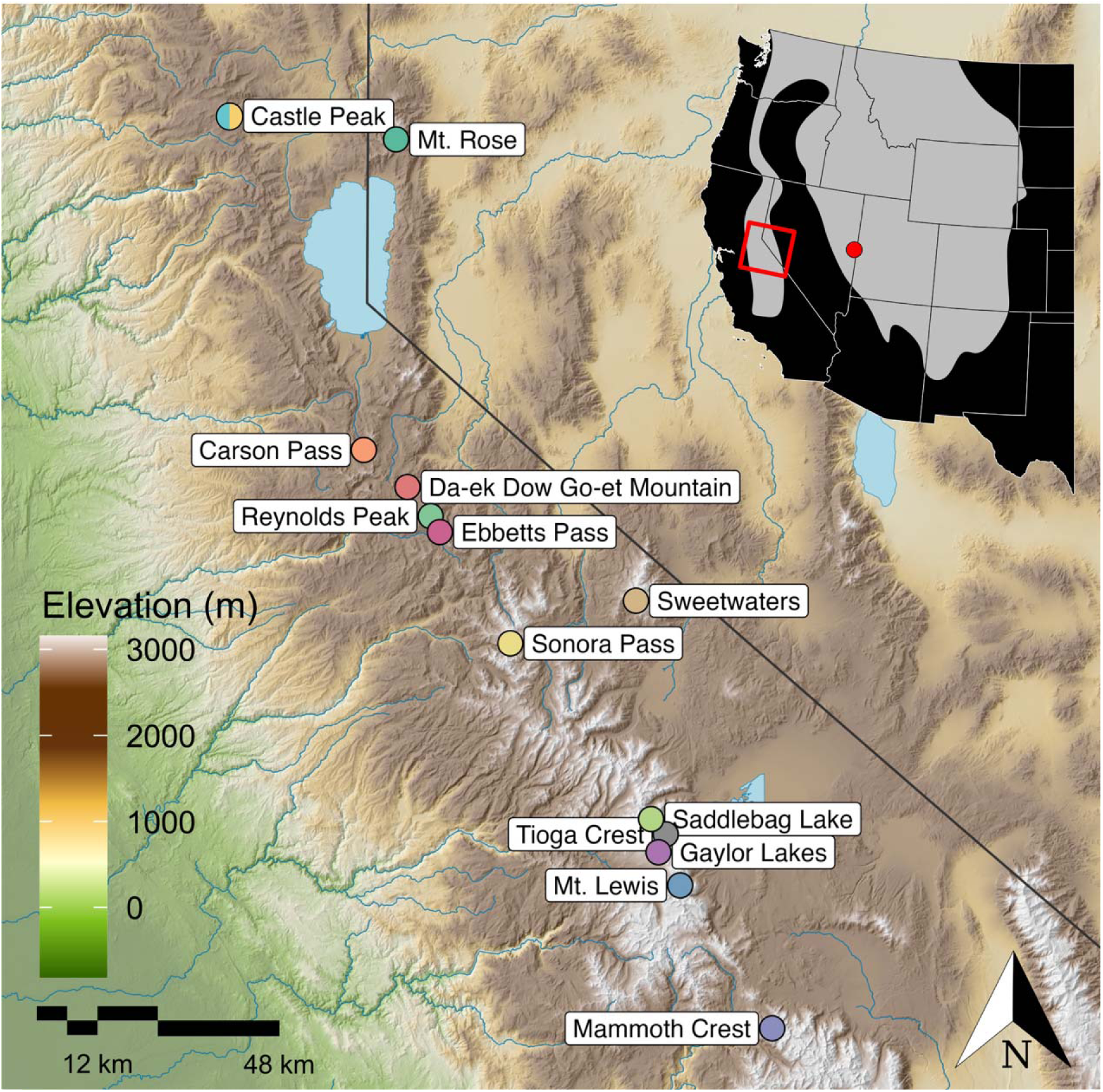
Map of 13 collection localities sampled within the Sierra Nevada mountains in the Western United States between 1991 and 1995. Castle Peak is labeled with a multicolored point to indicate hypothesized sympatric populations. Labels are directly horizontal to the point they refer to. The inset map shows the western U.S. distribution of *Oeneis chryxus* in gray and the study area in the Sierra Nevada Mountains of California, USA, outlined in red. One additional locality was sampled in the Snake Mountains of eastern Nevada for use as an outgroup (indicated by a red point on the inset map).

In 2024, we extracted DNA from the homogenates using the DNeasy Blood and Tissue Kit (Qiagen Inc., Alameda, CA, USA). We then built a reduced representation genomic library for 511 individuals across the Sierra, including 98 from Castle Peak, using previously described methodologies (Parchman et al. 2012, Gompert et al. 2014). DNA was digested with the EcoR1 and Mse1 restriction enzymes and Illumina adaptors with unique 8–10bp individual multiplex identifier (MID) sequences were ligated to the resulting fragments. We then amplified fragments using two rounds of PCR using iProof high-fidelity polymerase (BioRad Inc.) and pooled the resulting amplicons. Fragments between 300 and 450bp were selected using a BluePippin (Sage Science Inc., Beverly, MA, USA), and the resulting fragments were sequenced on two lanes of an Illumina Nextseq 2000 (single read, 150bp) at the SUNY Upstate Medical University’s Molecular Analysis Core (SUNYMAC, Syracuse, NY).

### Bioinformatics

PhiX reads were removed by assembly to the PhiX genome using bowtie version 1.1.2 (Langmead et al. 2009). We then removed the MIDs, short reads, and any reads that contained Mse1 adapter sequence. The remaining reads were written into separate fastq files for each individual (median = 3,861,495 reads per individual). We screened out 79 additional individuals who received fewer than 800,000 total reads. After this, we were left with 432 individuals, including 13, 10, 16, 22, and 17 individuals from Castle Peak from the years 1991-1995, respectively. After screening, all remaining reads were then aligned to an *O. chryxus* genome (MacDonald et al. 2024) using the mem algorithm of bwa version 0.7.18 (Li and Durbin 2009).

After alignment, we performed variant calling and additional filtering twice: first using all individuals and then only those collected at Castle Peak. In both cases, variable sites were identified using bcftools version 1.9 (Li et al. 2009) using the mpileup and call commands, ignoring indels and retaining variable sites if the posterior probability that the nucleotide was invariant was < 0.05. We further filtered this resulting variant call format file using custom scripts where variable sites were removed if they had a sequence depth of less than twice the number of individuals, a mapping quality less than 30, an absolute value of the mapping quality rank sum test greater than 1.96, an absolute value of the read position rank sum test greater than 1.96, an absolute value of the base quality rank sum test greater than 1.96, minor allele frequency less than 0.05, or missing data for more than 50% of the individuals. We limited variable sites to one per contig to minimize linkage disequilibrium. These steps resulted in a final dataset of 432 individuals and 27,304 variable sites across all populations and 78 individuals and 21,673 variable sites when only looking at Castle Peak.

### Statistics

To understand the distribution of genetic variation between butterflies collected in odd versus even years at Castle Peak, we estimated genotypes, allele frequencies, and admixture proportions using the Bayesian admixture algorithm entropy (Gompert et al. 2014, Shastry et al. 2021). We ran this algorithm twice on both the full and Castle Peak-only datasets. For both analyses, we fit multiple models, each specifying a different number of hypothetical ancestral populations: 2-15 for the full dataset and 2-6 for the Castle Peak-only dataset. We also estimated inter-class ancestry for the Castle Peak-only dataset using entropy (k=2) with the ancestry-complement model. For each model run, two MCMC simulations of 105,000 steps with a burn-in of 5,000 were run, thinning to retain every 10^th^ step. We assessed model convergence by calculating the mean Gelman-Rubin convergence diagnostic for admixture proportions of each individual (Gelman and Rubin 1992, Brooks and Gelman 1998) for each chain with the package coda version 0.19–1 in R (Plummer et al. 2006, R Core Team 2023). We also saved the deviance information criterion for each model to use for model comparison.

We investigated evolutionary relationships among individuals by building approximate maximum-likelihood phylogenetic trees with 1000 bootstrap replicates using FASTREE v2.1.11 (Price et al. 2009). Trees were constructed by concatenating SNPs from the filtered variant call file generated from the full dataset. This methodology is not suitable for estimating divergence times; however, we were interested only in topology and make no inferential statements about the timing of population differentiation. The tree was visualized with the ggtree package in R (Yu et al. 2017).

Genetic diversity was estimated for each site-collection year combination by calculating expected heterozygosity, the Watterson estimator (θ_w_) (Watterson 1975), and nucleotide diversity (θ_π_) (Nei and Li 1979). The Watterson estimator and nucleotide diversity were estimated using ANGSD v0.940 (Korneliussen et al. 2014). For each metric, the average was calculated across individuals for each site-collection-year combination.

To quantify the magnitude of differentiation between collection sites, we calculated genome-average pairwise Nei’s G_ST_ (Nei 1973). This was calculated as the average across all loci and all pairwise site-collection-year combinations, calculated as (average H_t_ – H_s_) / (average H_t_). To examine the relationship between G_ST_ and geographic distance, we used a subset of the full G_ST_ matrix by removing all even-year G_ST_ comparisons for sites with sampling in both even and odd years. By removing even-year samples, we compared genetic distances across space alone, removing any additional differentiation due to time. We then linearized values using G_ST_ / (1 – G_ST_) and performed a robust linear regression after using the MASS package (Venables and Ripley 2002) to quantify the relationship between G_ST_ and distance, which we used only to obtain an effect size, not for hypothesis testing. To infer differences between sites due to spatial and temporal isolation, we generated confidence intervals around each population’s G_ST_ using 1000 bootstrap samples and report the mean and 95% CI for each pairwise comparison. For differences over distance, Castle Peak odd years were used as the comparator group, and we examined bootstrapped pairwise G_ST_ values between Castle Peak odd-year samples and all other collection locations. To examine temporal differences, we compared bootstrapped pairwise G_ST_ values for samples collected from the same site in different odd years, different even years, or between odd and even years. All analyses were conducted in R (R Core Team 2023).

## Results

Sequencing produced 1,988,942,122 reads with MIDs for 511 individuals (median = 3,861,495 reads per individual, minimum = 786, maximum = 8,786,952). For analysis of the full dataset (from all collection sites), 191,225,644 reads were retained in the final assembly after filtering, for a SNP set of 27,034 loci. Mean sequence depth was 17.3 reads per individual per locus (standard deviation = 13.7). For analysis of the Castle Peak-only dataset, 16,401,409 reads were retained after filtering, yielding a SNP set of 21,673 loci. Mean sequence depth was 19.7 reads per individual per locus (standard deviation = 11.8). All models for the Castle Peak-only dataset converged; however, for the full dataset, only models with 10 or fewer clusters converged, and only results from these were implemented into further analyses (Table S1).

Principal components analysis of the posterior genotype estimates from the Castle Peak-only dataset revealed clear separation between individuals collected in odd and even years (Fig. 2A). Differences between years accounted for the greatest axis of variation (PC1, 7% of the variation; Fig. 2A). This separation is further supported by results from the admixture analysis, which shows that regardless of the number of hypothesized ancestral populations (k = 2-5 had comparable DIC scores, see Fig. S1), patterns of admixture remain the same. Odd- and even-year individuals are largely distinct, but a few individuals caught in even years show evidence of hybrid ancestry (Fig. 2B, Fig. S2). This is further explored in the results of the ancestry complement model (Fig. 2C), which indicates that one of these individuals, with approximately 0.5 admixture proportion from each population and 0.5 interclass ancestry, represents an F_2_ hybrid between even and odd year cohorts, while the other hybrid is a later generation backcross (Fig. 2C).

**Figure 2.**
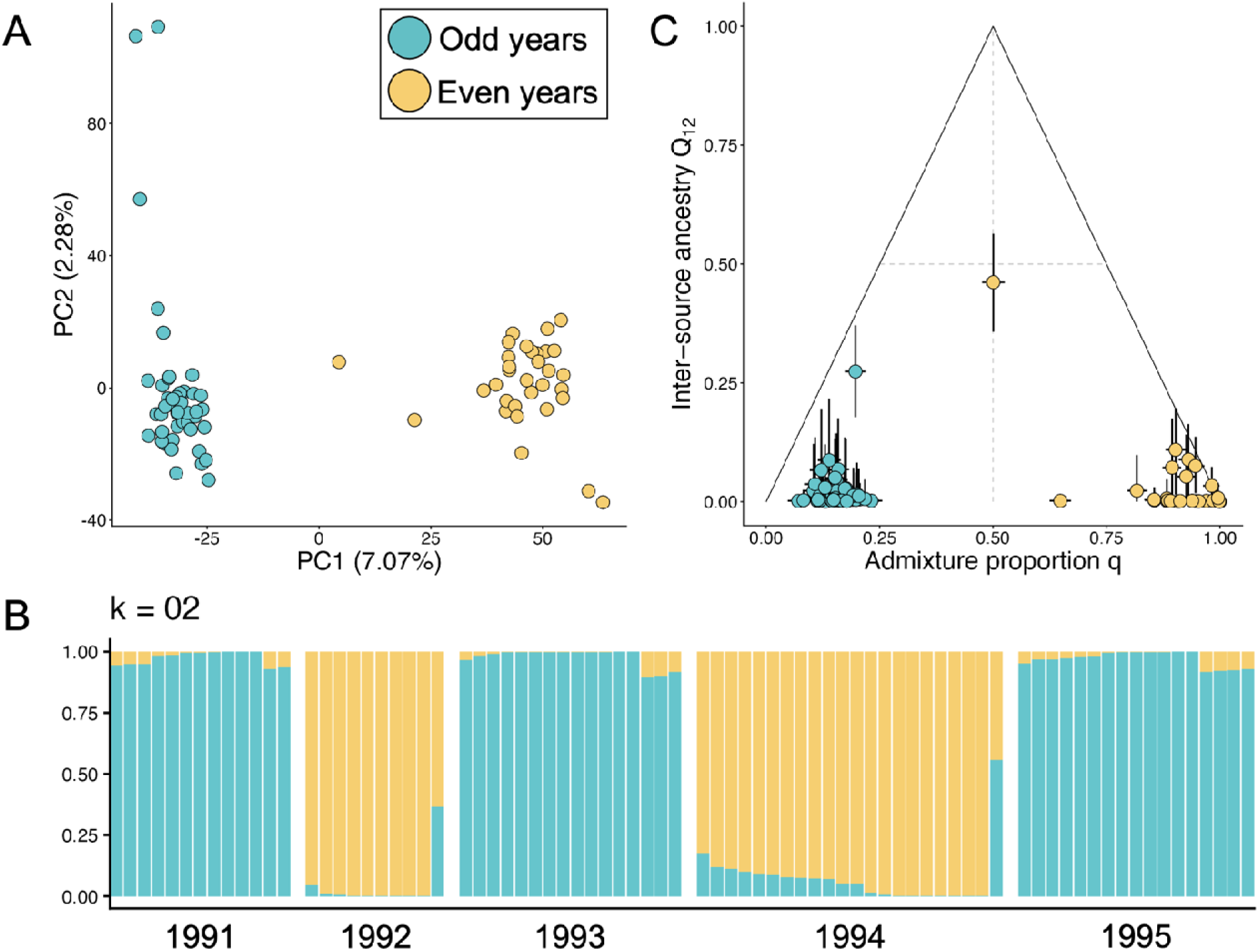
The population structure of *Oeneis chryxus* collected from Castle Peak between 1991 and 1995. A) Principal component analysis of posterior genotype probabilities summarizing variation across 78 individuals using 21,673 loci. B) Admixture proportions from Entropy models with two hypothesized ancestral populations (k=2) organized by collection year (results for k=3-6 can be found in Fig. S2). C) Results from the ancestry complement model in Entropy, where the x-axis is genome-average ancestry and the y-axis is the proportion of sampled loci that are heterozygous for ancestry.

We then examined the relatedness of the hypothesized populations by constructing a phylogenetic tree using the complete dataset from all 13 collection sites, which represents genetic variation within the Sierra Nevada, and an additional outgroup from the Great Basin (Fig. 3; see Fig. S3 for the full tree). The tree topology shows that even-year individuals are derived from odd-year individuals, the odd-year individuals are paraphyletic with respect to the even-year individuals, and this most likely occurred in a single event rather than a continuous series of founding events. We also found that the average genetic diversity across all metrics for individuals collected in even years is slightly lower than in odd years, suggesting a legacy of a founding event, although the differences are small relative to the overall variability (Table S2). Taken together, these results support the hypothesis that the odd- and even-year populations at Castle Peak are at least partially reproductively isolated in time, originated from a single event, and occasionally exchange individuals.

**Figure 3.**
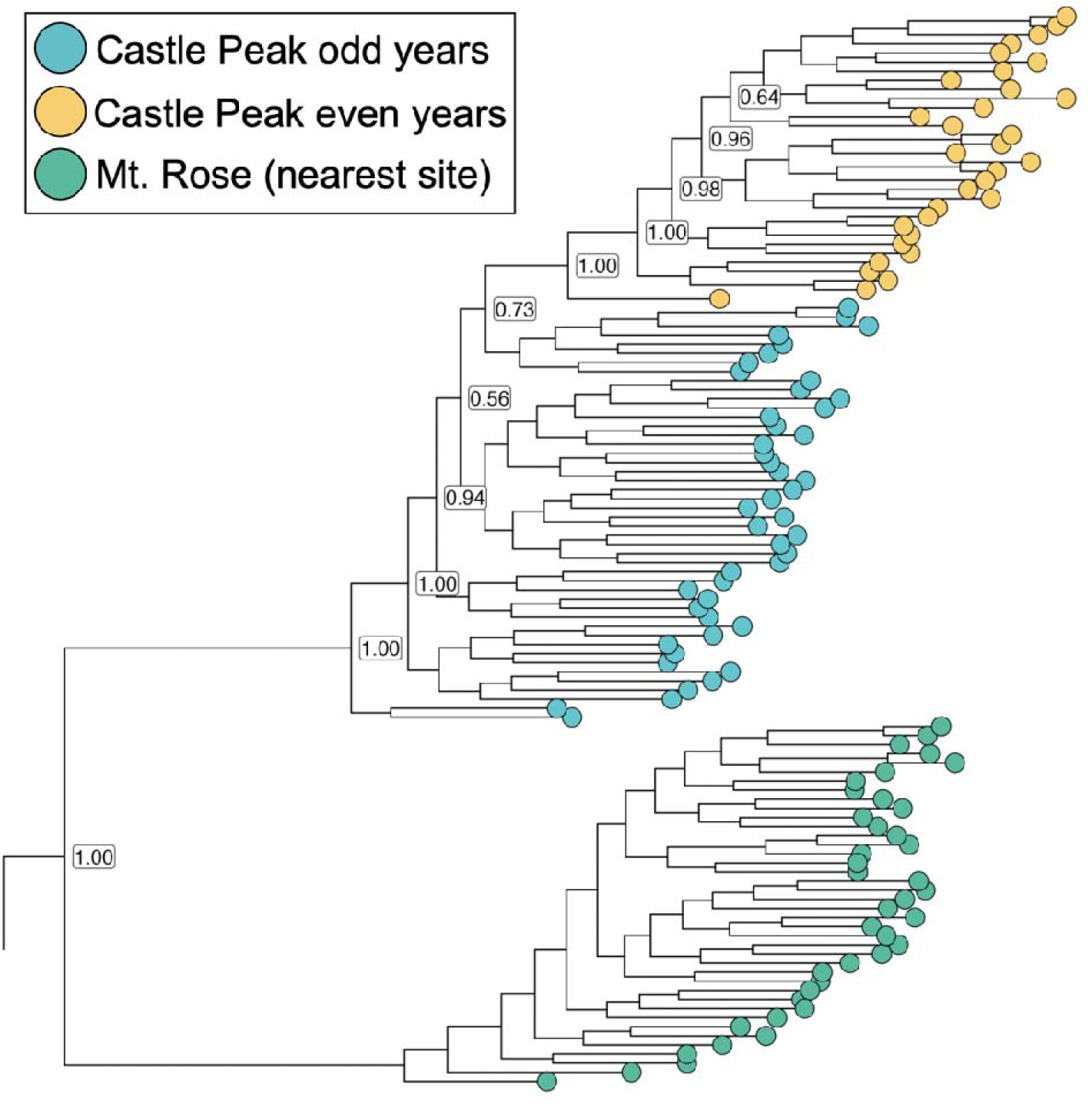
Subset of phylogenetic tree generated from all 14 source populations, showing Castle Peak and the closest geographic site (the full tree can be found in Fig. S4). Important nodes for establishing the topology are displayed and labeled with proportional bootstrap values from 1000 replicates.

After establishing that odd- and even-year cohorts are genetically distinguishable populations, we quantified the magnitude of differentiation. We did this using pairwise G_ST_ values observed across the full dataset (Table S3, Table S4, Table S5). We observed a strong positive relationship between pairwise distance and G_ST_, with every 0.1 change in G_ST_ associated with approximately 52 km (se ± 1km) (Fig. 4A, Table S6). We found that all populations, across space and time, differed from the Castle Peak odd years. While the largest differences were observed between sites that were more spatially distant, we also found differences between temporally separated populations. Specifically, even and odd years at Castle Peak yield an average G_ST_ difference of 0.048 (Bootstrap CI: 0.044-0.057) (Fig. 4B, Table S3). When translated into geographic distance, the average distance between odd and even year populations at Castle Peak is approximately equivalent to a geographic distance of 26km. While a limited sample size prevented complete analyses, we also found that the three individuals collected from the Sweetwater mountains in 1992 differed in pairwise G_ST_ by 0.048 (Bootstrap CI: 0.048-0.049) and 0.052 (Bootstrap CI: 0.051-0.053) from individuals collected in 1993 and 1995, respectively (Table S4).

**Figure 4.**
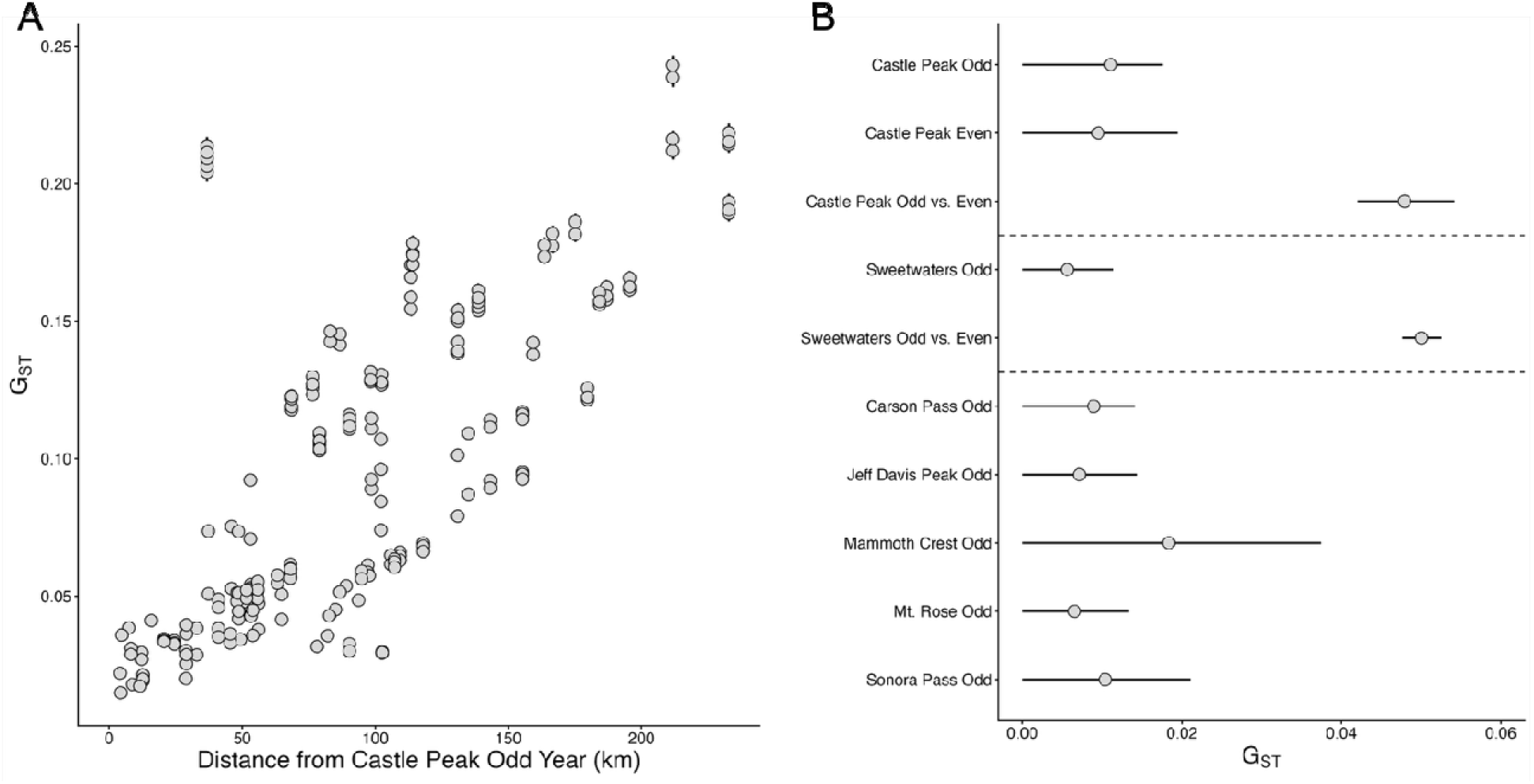
G_ST_ values compared across space and time. A) G_ST_ values across the 13 collection localities in the Sierra Nevada mountains compared to each of the Castle Peak odd years. B) Within-site G_ST_ values from sites where multiple samples were taken. Odd indicates that odd years are being compared with other odd years, while odd vs. even indicates that odd years are being compared to even years (from the same site). The confidence intervals around each G_ST_ estimate were generated using 1000 bootstrap samples.

## Discussion

Population divergence due to geographic separation is a classic prediction of evolutionary theory (Wright 1943); however, in specific systems, a similar pattern can arise from temporal separation (Hendry and Day 2005). In some cases, allochrony and selection have led to substantial divergence (Taylor and Friesen 2017), but the frequency and magnitude at which drift alone can lead to temporally structured populations across years remain open questions (Devaux and Lande 2008). Here, we examined whether allochronic sympatry occurred in the butterfly *Oeneis chryxus ivallda*, a species that develops over two years but occurs in certain locations each year. We found that individuals collected over five consecutive years at Castle Peak, CA, exhibited clear population structure between odd- and even-year cohorts, with minimal admixture. The observed temporal separation was comparable to 26 km in space. These results provide a clear example of allochronic separation in an extreme environment and raise questions about whether similar processes may occur elsewhere.

Reproductive isolation by time is related to both the temporal distance (relative to the length of the organism’s reproductive stage) and gene flow between different cohorts (Hendry and Day 2005). Allochronic separation across years has previously been observed in organisms with rigid reproductive cycle timing and where cross-cohort breeding is rare, such as pink salmon (*Oncorhynchus gorbuscha*) and lousewort (*Pedicularis hallaisanensis*) (Aspinwall 1974, Kim et al. 2024). In contrast, although *O. chryxus* typically overwinters twice, it is not obligate, as rearing has shown that both diapauses can be prevented under specific conditions (James and Nunnallee 2011). Yet, although individuals are not physiologically prevented from emerging in the alternate year, such events appear to be relatively uncommon, as we found only two individuals with evidence of hybrid ancestry among the 78 sequenced individuals. Given this, it seems likely that the environmental conditions necessary to compress development into a single year or extend it to three years were, at the very least, unusual. Since *O. chryxus* inhabits montane environments, the magnitude of gene flow may depend on local microclimatic heterogeneity, with greater within-site variability increasing the probability that at least some individuals switch to alternate allochronic populations (Greiser et al. 2022). Castle Peak comprises a single ridgeline and is smaller and less heterogeneous than the vast alpine landscapes that characterize much of *O. chryxus’* range to the south. In this relatively simple landscape, reduced instances of individuals switching would increase the strength of drift between the two sympatric populations. Regardless of the mechanisms underlying admixture, the *O. chryxus* populations at Castle Peak represent a clear example of how allochronic genetic differentiation can persist despite minor gene flow.

Although some contact between the two allochronic populations indicates that individuals occasionally switch years, only one site in the Sierra Nevada has been known to host a large even-year population, suggesting additional barriers to the founding of new populations. An explanation may be found in the severe Allee effects that founders would face (Courchamp et al. 1999), including low genetic diversity and trouble finding mates, which can be especially important in insects (Kramer et al. 2009). Overcoming this would be more likely if many individuals switched years at the same time, making environments where this was more probable more likely to host two co-occurring populations. Castle Peak was both the northernmost known *O. chryxus* population in the Sierra Nevada and among the lowest elevation, and as a result, has a comparatively longer growing season. It may be that incipient off-year populations arise more frequently at places like this, at range edges, where exposure to more extreme conditions (relative to the core range) increases developmental variation and, consequently, the likelihood of switching events. Given that the even-year population is derived from the odd-year population and that large-scale climate events shape synchrony across Northern California butterflies (Pardikes et al. 2017), it is plausible that the even-year population originated from a single extreme climate event. Of course, Castle Peak is only a single location, and the possibility remains that the founding event was an anomaly and that distribution of where off-year populations establish and persist may be random, or at least appear to be, even if abiotic and biotic processes are acting at fine scales.

The *Oeneis chryxus ivallda* populations at Castle Peak illustrate a case in which selection is unlikely to be driving differentiation, because individuals from both cohorts ultimately experience the same environment, unlike in systems with seasonal allochrony, where distinct environments across time can promote divergence (Taylor and Friesen 2017). Yet our results show that separation across years, acting through stochastic processes alone, can generate population structure comparable to that produced over short geographic distances. While the equally sized even and odd year populations of Castle Peak may indeed have been the exception across the range, the few even year individuals sequenced from the Sweetwater mountains also exhibited similar differentiation from the sympatric odd year cohort (Fig. 4). The butterflies at that site presented the pattern more typical of *O. chryxus* across much of the Sierra Nevada: a large, odd year cohort with infrequently observed even year individuals (MacDonald et al. 2024). Given the observed differentiation, it remains possible that smaller but still differentiated populations of *O. chryxus* may be widespread, as demonstrated in other insects (Heliövaara et al. 1988). Further still, while this and similar work have focused on cold settings (Heliövaara et al. 1988, Vila and Björklund 2004, Gradish et al. 2015, 2019), prolonged development across years is a feature of many extreme environments, including those in temperate hot and dry regions as well as in the tropics (Danks 2013). Even within the *Oeneis* genus, other lower-elevation species are believed to have biennial development (Scott 1986), and such systems may provide key insights into the frequency and variation of yearly allochronic divergence in extreme environments.

## Author Contributions

C.A.H.: Methodology, Formal analysis, Investigation, Data Curation, Writing - Original Draft, Visualization; C.C.N. Conceptualization, Methodology, Investigation, Data Curation, Writing - Review & Editing, Supervision; K.L.B.: Methodology, Writing - Review & Editing; J.A.F.: Methodology, Writing - Review & Editing; M.L.F.: Writing - Review & Editing; G.G.C.: Writing - Review & Editing; A.M.S.: Writing - Review & Editing; E.M.G: Writing - Review & Editing, Supervision, Funding acquisition.

## Acknowledgments

We want to thank J. Van Wangtendonk and the Yosemite field station for help acquiring collecting permits, G. Kareofelas for collecting the Sonora Pass 1989 population sample, S. Graves and K. Roberg for assistance with collecting, and J. Mori for important information on collecting localities and natural history. We also thank Trevor Faske for providing feedback on a previous version of this manuscript.

## References

Aspinwall, N. 1974. Genetic Analysis of North American Populations Of The Pink Salmon, Oncorhynchus Gorbuscha, Possible Evidence for the Neutral Mutation-Random Drift Hypothesis. Evolution; International Journal of Organic Evolution 28:295–305.

Bell, K. L., C. A. Hamm, A. M. Shapiro, and C. C. Nice. 2017. Sympatric, temporally isolated populations of the pine white butterfly Neophasia menapia, are morphologically and genetically differentiated. PLOS ONE 12:e0176989.

Berlocher, S. H., and J. L. Feder. 2002. Sympatric Speciation in Phytophagous Insects: Moving Beyond Controversy? Annual Review of Entomology 47:773–815.

Bohonak, A. J. 1999. Dispersal, Gene Flow, and Population Structure. The Quarterly Review of Biology 74:21–45.

Brooks, S. P., and A. Gelman. 1998. General Methods for Monitoring Convergence of Iterative Simulations. Journal of Computational and Graphical Statistics 7:434–455.

Courchamp, F., T. Clutton-Brock, B. Grenfell, F. Courchamp, T. Clutton-Brock, B. Grenfell, F. Courchamp, T. Clutton-Brock, B. Grenfell, F. Courchamp, T. Clutton-Brock, B. Grenfell, F. Courchamp, T. Clutton-Brock, and B. Grenfell. 1999. Inverse density dependence and the Allee effect. Trends in Ecology & Evolution 14:405–410.

Danks, H. V. 1992. Long Life Cycles In Insects. The Canadian Entomologist 124:167–187.

Danks, H. V. 2013. The range of insect dormancy responses. EJE 99:127–142.

Devaux, C., and R. Lande. 2008. Incipient allochronic speciation due to non-selective assortative mating by flowering time, mutation and genetic drift. Proceedings of the Royal Society B: Biological Sciences 275:2723–2732.

Feder, J. L., C. A. Chilcote, and G. L. Bush. 1988. Genetic differentiation between sympatric host races of the apple maggot fly Rhagoletis pomonella. Nature 336:61–64.

Friesen, V. L., A. L. Smith, E. Gómez-Díaz, M. Bolton, R. W. Furness, J. González-Solís, and L. R. Monteiro. 2007. Sympatric speciation by allochrony in a seabird. Proceedings of the National Academy of Sciences 104:18589–18594.

Fudickar, A. M., T. J. Greives, J. W. Atwell, C. A. Stricker, and E. D. Ketterson. 2016. Reproductive Allochrony in Seasonally Sympatric Populations Maintained by Differential Response to Photoperiod: Implications for Population Divergence and Response to Climate Change. The American Naturalist 187:436–446.

Fukami, H., M. Omori, K. Shimoike, T. Hayashibara, and M. Hatta. 2003. Ecological and genetic aspects of reproductive isolation by different spawning times in Acropora corals. Marine Biology 142:679–684.

Gelman, A., and D. B. Rubin. 1992. Inference from Iterative Simulation Using Multiple Sequences. Statistical Science 7:457–472.

Gompert, Z., L. K. Lucas, C. A. Buerkle, M. L. Forister, J. A. Fordyce, and C. C. Nice. 2014. Admixture and the organization of genetic diversity in a butterfly species complex revealed through common and rare genetic variants. Molecular Ecology 23:4555–4573.

Gradish, A. E., N. Keyghobadi, and G. W. Otis. 2015. Population genetic structure and genetic diversity of the threatened White Mountain arctic butterfly (Oeneis melissa semidea). Conservation Genetics 16:1253–1264.

Gradish, A. E., N. Keyghobadi, F. A. H. Sperling, and G. W. Otis. 2019. Population genetic structure and assessment of allochronic divergence in the Macoun’s Arctic (Oeneis macounii) butterfly. Canadian Journal of Zoology 97:121–130.

Greiser, C., L. von Schmalensee, O. Lindestad, K. Gotthard, and P. Lehmann. 2022. Microclimatic variation affects developmental phenology, synchrony and voltinism in an insect population. Functional Ecology 36:3036–3048.

Halsch, C. A., A. M. Shapiro, J. H. Thorne, K. C. Rodman, A. Parra, L. A. Dyer, Z. Gompert, A. M. Smilanich, and M. L. Forister. 2024. Thirty-six years of butterfly monitoring, snow cover, and plant productivity reveal negative impacts of warmer winters and increased productivity on montane species. Global Change Biology 30:e17044.

Heliövaara, K., and R. Väisänen. 1987. Geographic variation in the life-history of Aradus cinnamomeus and a breakdown mechanism of the reproductive isolation of allochronic bugs (Heteroptera, Aradidae). Annales Zoologici Fennici 24:1–17.

Heliövaara, K., R. Väisänen, J. Hantula, J. Lokki, and A. Saura. 1988. Genetic differentiation in sympatric but temporally isolated pine bark bugs, Aradus cinnamomeus (Heteroptera). Hereditas 109:29–36.

Heliövaara, K., R. Väisänen, and C. Simon. 1994. Evolutionary ecology of periodical insects. Trends in Ecology & Evolution 9:475–480.

Hendry, A. P., and T. Day. 2005. Population structure attributable to reproductive time: isolation by time and adaptation by time. Molecular Ecology 14:901–916.

Hovanitz, W. 1940. Ecology Color Variation in a Butterfly and the Problem of “Protective Coloration.” Ecology 21:371–380.

Hutchings, J. A., and M. E. Jones. 1998. Life history variation and growth rate thresholds for maturity in Atlantic salmon, Salmo salar. Canadian Journal of Fisheries and Aquatic Sciences 55:22–47.

Ito, H., S. Kakishima, T. Uehara, S. Morita, T. Koyama, T. Sota, J. R. Cooley, and J. Yoshimura. 2015. Evolution of periodicity in periodical cicadas. Scientific Reports 5:14094.

James, D. G., and D. Nunnallee. 2011. Life Histories of Cascadia Butterflies. Oregon State University Press, Corvallis, OR.

Johnston, S. E., P. Orell, V. L. Pritchard, M. P. Kent, S. Lien, E. Niemelä, J. Erkinaro, and C. R. Primmer. 2014. Genome-wide SNP analysis reveals a genetic basis for sea-age variation in a wild population of Atlantic salmon (Salmo salar). Molecular Ecology 23:3452–3468.

Kim, S., B.-D. Lee, C. W. Lee, H.-J. Park, J. E. Hwang, H. B. Park, Y.-J. Kim, D. Jeon, and Y.-J. Yoon. 2024. Strict biennial lifecycle and anthropogenic interventions affect temporal genetic differentiation in the endangered endemic plant, Pedicularis hallaisanensis. Frontiers in Plant Science 15.

Koenig, W. D., and A. M. Liebhold. 2013. Avian Predation Pressure as a Potential Driver of Periodical Cicada Cycle Length. The American Naturalist 181:145–149.

Korneliussen, T. S., A. Albrechtsen, and R. Nielsen. 2014. ANGSD: Analysis of Next Generation Sequencing Data. BMC Bioinformatics 15:356.

Kramer, A. M., B. Dennis, A. M. Liebhold, and J. M. Drake. 2009. The evidence for Allee effects. Population Ecology 51:341.

Langmead, B., C. Trapnell, M. Pop, and S. L. Salzberg. 2009. Ultrafast and memory-efficient alignment of short DNA sequences to the human genome. Genome Biology 10:R25.

Li, H., and R. Durbin. 2009. Fast and accurate short read alignment with Burrows-Wheeler transform. Bioinformatics (Oxford, England) 25:1754–1760.

Li, H., B. Handsaker, A. Wysoker, T. Fennell, J. Ruan, N. Homer, G. Marth, G. Abecasis, and R. Durbin. 2009. The Sequence Alignment/Map format and SAMtools. Bioinformatics 25:2078–2079.

MacDonald, Z. G., S. Schoville, M. Escalona, M. P. A. Marimuthu, O. Nguyen, N. Chumchim, C. W. Fairbairn, W. Seligmann, E. Toffelmier, T. Gillespie, and H. B. Shaffer. 2024. A genome assembly for the Chryxus Arctic (Oeneis chryxus), the highest butterfly in North America. Journal of Heredity:esae051.

Nei, M. 1973. Analysis of gene diversity in subdivided populations. Proceedings of the National Academy of Sciences of the United States of America 70:3321–3323.

Nei, M., and W. H. Li. 1979. Mathematical model for studying genetic variation in terms of restriction endonucleases. Proceedings of the National Academy of Sciences of the United States of America 76:5269–5273.

Nice, C. C., and A. M. Shapiro. 2001. Patterns of Morphological, Biochemical, and Molecular Evolution in the *Oeneis chryxus* Complex (Lepidoptera: Satyridae): A Test of Historical Biogeographical Hypotheses. Molecular Phylogenetics and Evolution 20:111–123.

Parchman, T. L., Z. Gompert, J. Mudge, F. D. Schilkey, C. W. Benkman, and C. A. Buerkle. 2012. Genome-wide association genetics of an adaptive trait in lodgepole pine. Molecular Ecology 21:2991–3005.

Pardikes, N. A., J. G. Harrison, A. M. Shapiro, and M. L. Forister. 2017. Synchronous population dynamics in California butterflies explained by climatic forcing. Royal Society Open Science 4:170190.

Peterson, B. K., J. N. Weber, E. H. Kay, H. S. Fisher, and H. E. Hoekstra. 2012. Double Digest RADseq: An Inexpensive Method for De Novo SNP Discovery and Genotyping in Model and Non-Model Species. PLOS ONE 7:e37135.

Plummer, M., N. Best, K. Cowles, and K. Vines. 2006. CODA: Convergence Diagnosis and Output Analysis for MCMC. R News 6:7–11.

Porter, A. L., and A. M. Shapiro. 1989. Genetics and Biogeograph7 of the Oeneis chryxus Complex (Satyrinae) in California. Journal of Research on Lepidoptera 28:263–276.

Price, M. N., P. S. Dehal, and A. P. Arkin. 2009. FastTree: Computing Large Minimum Evolution Trees with Profiles instead of a Distance Matrix. Molecular Biology and Evolution 26:1641–1650.

R Core Team. 2023. R: A Language and Environment for Statistical Computing. R Foundation for Statistical Computing, Vienna, Austria.

Ritchie, M. G. 2001. Chronic speciation in periodical cicadas. Trends in Ecology & Evolution 16:59–61.

Rousset, F. 1997. Genetic differentiation and estimation of gene flow from F-statistics under isolation by distance. Genetics 145:1219–1228.

Rundle, H. D., and P. Nosil. 2005. Ecological speciation. Ecology Letters 8:336–352.

Santos, H., J. Rousselet, E. Magnoux, M.-R. Paiva, M. Branco, and C. Kerdelhué. 2007. Genetic isolation through time: allochronic differentiation of a phenologically atypical population of the pine processionary moth. Proceedings of the Royal Society B: Biological Sciences 274:935–941.

Scott, J. A. 1986. The Butterflies of North America: A Natural History and Field Guide. Stanford University Press, Stanford.

Shastry, V., P. E. Adams, D. Lindtke, E. G. Mandeville, T. L. Parchman, Z. Gompert, and C. A. Buerkle. 2021. Model-based genotype and ancestry estimation for potential hybrids with mixed-ploidy. Molecular Ecology Resources 21:1434–1451.

Simon, C., J. R. Cooley, R. Karban, and T. Sota. 2022. Advances in the Evolution and Ecology of 13- and 17-Year Periodical Cicadas. Annual Review of Entomology 67:457–482.

Slatkin, M. 1987. Gene Flow and the Geographic Structure of Natural Populations. Science 236:787–792.

Sota, T., S. Yamamoto, J. R. Cooley, K. B. R. Hill, C. Simon, and J. Yoshimura. 2013. Independent divergence of 13- and 17-y life cycles among three periodical cicada lineages. Proceedings of the National Academy of Sciences 110:6919–6924.

Taylor, R. S., and V. L. Friesen. 2017. The role of allochrony in speciation. Molecular Ecology 26:3330–3342.

Venables, W. N., and B. D. Ripley. 2002. Modern Applied Statistics with S. Fourth. Springer, New York.

Vila, M., and M. Björklund. 2004. Testing biennialism in the butterfly Erebia palarica (Nymphalidae: Satyrinae) by mtDNA sequencing. Insect Molecular Biology 13:213–217.

Watterson, G. A. 1975. On the number of segregating sites in genetical models without recombination. Theoretical Population Biology 7:256–276.

Wright, S. 1943. Isolation by Distance. Genetics 28:114–138.

Yu, G., D. K. Smith, H. Zhu, Y. Guan, and T. T.-Y. Lam. 2017. ggtree: an r package for visualization and annotation of phylogenetic trees with their covariates and other associated data. Methods in Ecology and Evolution 8:28–36.

