## supplement for "Allochronic isolation between sympatric populations of an alpine butterfly"

This file includes:

Figure S1. DIC values for Entropy models run on the A) full dataset and B) Castle-only dataset.

Figure S2. Admixture proportions for *Oeneis chryxus* collected from Castle Peak between 1991 and 1995.

Figure S3. Full phylogenetic tree fit using the full dataset.

Table S1. Convergence diagnostics for each Entropy model run.

Table S2. Population level genetic diversity of each sample year at Castle Peak, CA.

Table S3. Pairwise G_ST_ between collection years and sites.

Table S4. Pairwise G_ST_ between collection years and sites continued.

Table S5. Pairwise G_ST_ between collection years and sites continued.

Table S6. Summary statistics for linear model predicting G_ST_ using geographic distance.

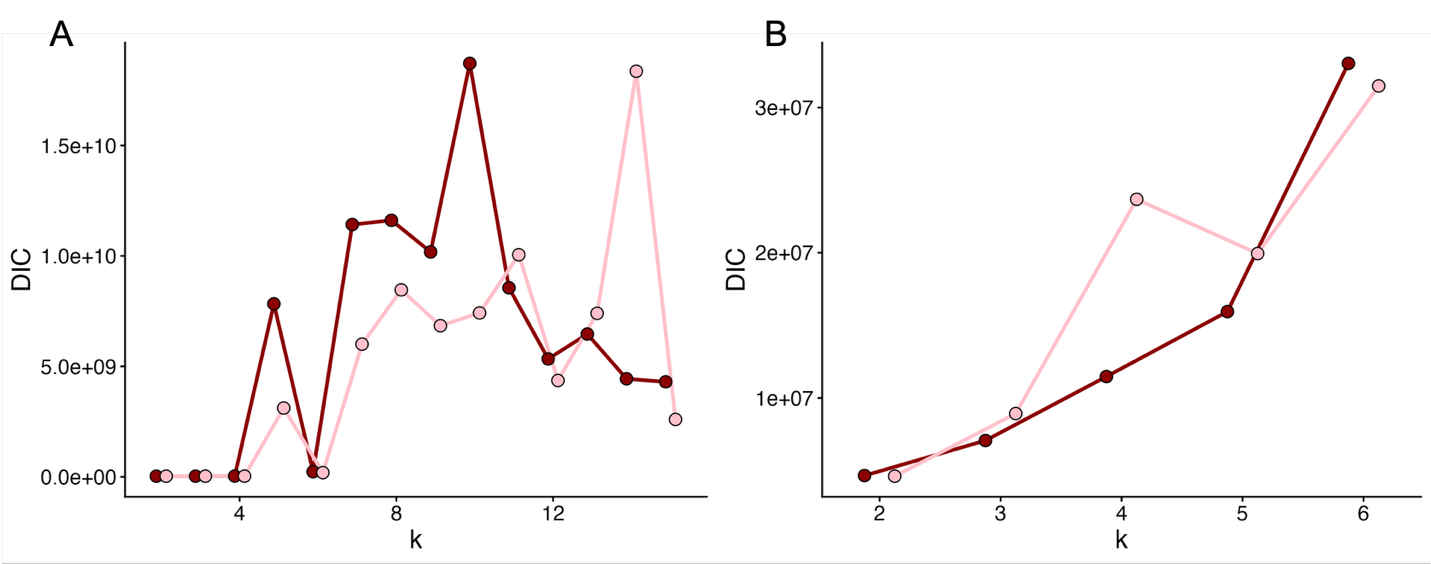

Fig. S1. DIC values for Entropy models run on the A) full dataset and B) Castle-only dataset. K is the number of hypothesized ancestral populations in the model, and the colors show different chains.

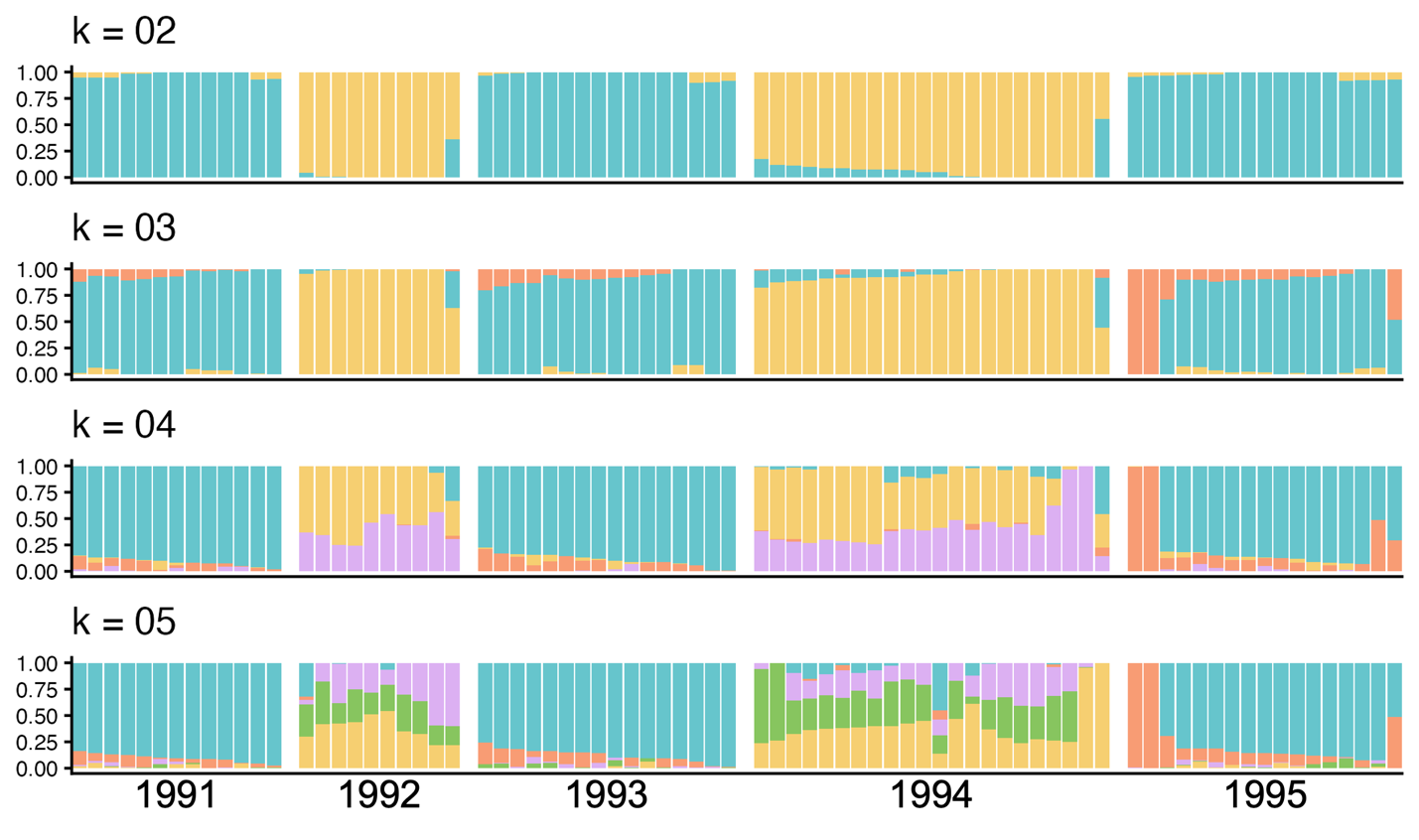

Figure S2. Admixture proportions for *Oeneis chryxus ivallda* collected from Castle Peak between 1991 and 1995. Results from entropy models with 2-6 hypothesized ancestral populations (k=2-6) organized by collection year.

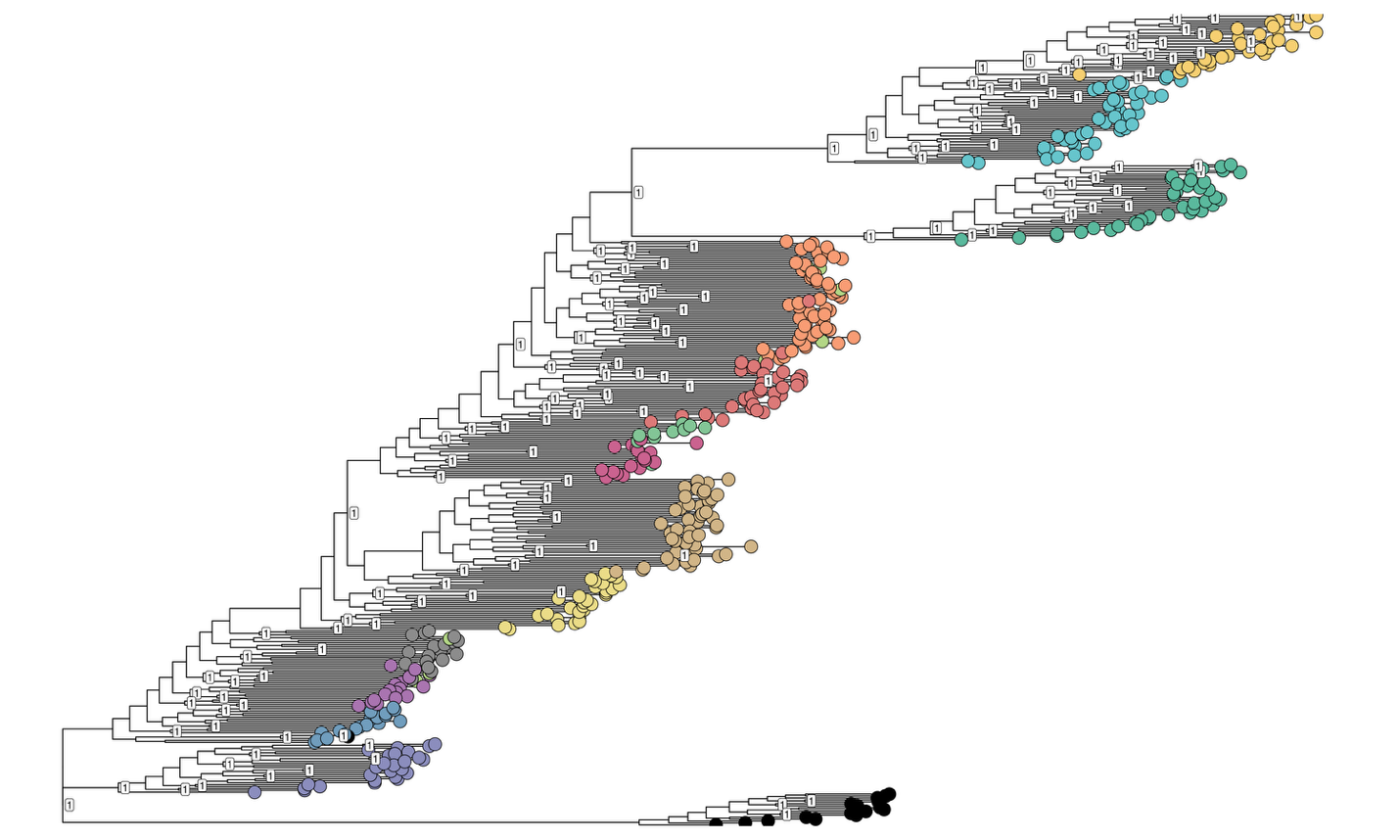

Figure S3. Full phylogenetic tree fit using the full dataset. Colors match those in Figure 1, and the black outgroup corresponds to the eastern population used as an outgroup.

| Table S1. Convergence diagnostics for each Entropy model run. Values show the mean Gelman Rubin statistic for individual parameters in the model. Models where k>10 were discarded from analysis of the full dataset. | | |
| --- | --- | --- |
| Number of clusters | Rhat from all data model | Rhat from Castle Peak only model |
| 2 | 1.01 | 1.05 |
| 3 | 1.01 | 1.04 |
| 4 | 1.02 | 1.04 |
| 5 | 1.02 | 1.08 |
| 6 | 1.05 |  |
| 7 | 1.03 |  |
| 8 | 1.03 |  |
| 9 | 1.05 |  |
| 10 | 1.05 |  |
| 11 | 1.68 |  |
| 12 | 1.05 |  |
| 13 | 1.41 |  |
| 14 | 1.95 |  |
| 15 | 1.43 |  |

| Table S2. Population level genetic diversity of each sample year at Castle Peak, CA. Standard deviations are shown in parentheses. | | | |
| --- | --- | --- | --- |
| **Collection year** | **Heterozygosity** | **Nucleotide diversity** | **Watterson theta** |
| 1991 | 0.202 | 0.00098 (0.00027) | 0.00093 (0.00021) |
| 1992 | 0.177 | 0.00086 (0.00026) | 0.00081 (0.00021) |
| 1993 | 0.205 | 0.00101 (0.00026) | 0.00099 (0.00020) |
| 1994 | 0.185 | 0.00090 (0.00023) | 0.00088 (0.00017) |
| 1995 | 0.203 | 0.00097 (0.00026) | 0.00095 (0.00020) |

| Table S3. Pairwise G_ST_ between collection years and sites. Observed G_ST_ values are show below the diagonal while bootstrapped 95% confidence intervals are shown above the diagonal. Continued in Table S3. | | | | | | | | |
| --- | --- | --- | --- | --- | --- | --- | --- | --- |
|  | **Castle 1991** | **Castle 1992** | **Castle 1993** | **Castle 1994** | **Castle 1995** | **Carson 1991** | **Carson 1993** | **Carson 1995** |
| **Castle 1991** |  | (0.052-0.055) | (0.017-0.017) | (0.044-0.045) | (0.017-0.018) | (0.107-0.11) | (0.108-0.111) | (0.105-0.108) |
| **Castle 1992** | 0.054 |  | (0.051-0.053) | (0.019-0.02) | (0.051-0.053) | (0.138-0.142) | (0.139-0.143) | (0.135-0.14) |
| **Castle 1993** | 0.017 | 0.052 |  | (0.042-0.044) | (0.015-0.016) | (0.103-0.107) | (0.104-0.108) | (0.101-0.105) |
| **Castle 1994** | 0.044 | 0.019 | 0.043 |  | (0.042-0.044) | (0.129-0.134) | (0.13-0.134) | (0.126-0.131) |
| **Castle 1995** | 0.017 | 0.052 | 0.015 | 0.043 |  | (0.104-0.108) | (0.105-0.108) | (0.102-0.105) |
| **Carson 1991** | 0.109 | 0.14 | 0.105 | 0.131 | 0.106 |  | (0.012-0.013) | (0.014-0.014) |
| **Carson 1993** | 0.109 | 0.141 | 0.106 | 0.132 | 0.107 | 0.013 |  | (0.013-0.014) |
| **Carson 1995** | 0.107 | 0.137 | 0.103 | 0.129 | 0.103 | 0.014 | 0.014 |  |
| **Ebbetts 1995** | 0.131 | 0.161 | 0.127 | 0.153 | 0.128 | 0.034 | 0.033 | 0.033 |
| **Gaylor 1995** | 0.162 | 0.194 | 0.158 | 0.185 | 0.159 | 0.066 | 0.065 | 0.063 |
| **Da ek dow go et 1993** | 0.116 | 0.148 | 0.112 | 0.139 | 0.113 | 0.021 | 0.021 | 0.021 |
| **Da ek dow go et 1995** | 0.115 | 0.146 | 0.111 | 0.138 | 0.112 | 0.02 | 0.019 | 0.02 |
| **Mammoth 1993** | 0.193 | 0.226 | 0.189 | 0.217 | 0.19 | 0.095 | 0.094 | 0.093 |
| **Mammoth 1995** | 0.218 | 0.253 | 0.214 | 0.243 | 0.215 | 0.117 | 0.116 | 0.114 |
| **Rose 1995** | 0.209 | 0.243 | 0.204 | 0.234 | 0.206 | 0.119 | 0.118 | 0.119 |
| **Rose 1993** | 0.214 | 0.249 | 0.209 | 0.239 | 0.211 | 0.123 | 0.122 | 0.123 |
| **Moriah 1995** | 0.166 | 0.198 | 0.161 | 0.189 | 0.162 | 0.069 | 0.068 | 0.066 |
| **Reynolds 1993** | 0.132 | 0.163 | 0.128 | 0.155 | 0.129 | 0.035 | 0.034 | 0.033 |
| **Saddlebag 1991** | 0.126 | 0.157 | 0.121 | 0.148 | 0.122 | 0.03 | 0.029 | 0.029 |
| **Sonora 1995** | 0.154 | 0.186 | 0.15 | 0.177 | 0.151 | 0.054 | 0.053 | 0.053 |
| **Sonora 1991** | 0.142 | 0.173 | 0.138 | 0.165 | 0.139 | 0.045 | 0.043 | 0.043 |
| **Sweetwaters 1995** | 0.2 | 0.232 | 0.196 | 0.223 | 0.197 | 0.097 | 0.095 | 0.094 |
| **Sweetwaters 1993** | 0.158 | 0.189 | 0.154 | 0.18 | 0.155 | 0.058 | 0.057 | 0.057 |
| **Sweetwaters 1992** | 0.161 | 0.193 | 0.157 | 0.184 | 0.158 | 0.062 | 0.06 | 0.06 |
| **Tioga 1995** | 0.16 | 0.193 | 0.156 | 0.184 | 0.157 | 0.063 | 0.062 | 0.06 |

| Table S4. Pairwise G_ST_ between collection years and sites. Observed G_ST_ values are show below the diagonal while bootstrapped 95% confidence intervals are shown above the diagonal. Continued in Table S4. | | | | | | | | |
| --- | --- | --- | --- | --- | --- | --- | --- | --- |
|  | **Ebbetts 1995** | **Gaylor 1995** | **Da ek dow go et 1993** | **Da ek dow go et 1995** | **Mammoth 1993** | **Mammoth 1995** | **Rose 1995** | **Rose 1993** |
| **Castle 1991** | (0.128-0.133) | (0.16-0.165) | (0.114-0.118) | (0.113-0.117) | (0.191-0.196) | (0.216-0.222) | (0.206-0.212) | (0.21-0.217) |
| **Castle 1992** | (0.159-0.164) | (0.192-0.197) | (0.145-0.15) | (0.144-0.149) | (0.223-0.229) | (0.249-0.256) | (0.24-0.247) | (0.245-0.252) |
| **Castle 1993** | (0.125-0.129) | (0.155-0.16) | (0.11-0.114) | (0.109-0.113) | (0.186-0.192) | (0.211-0.218) | (0.201-0.207) | (0.206-0.212) |
| **Castle 1994** | (0.151-0.155) | (0.183-0.188) | (0.136-0.141) | (0.135-0.14) | (0.214-0.22) | (0.24-0.247) | (0.231-0.238) | (0.236-0.243) |
| **Castle 1995** | (0.126-0.13) | (0.157-0.162) | (0.111-0.115) | (0.11-0.114) | (0.188-0.193) | (0.212-0.219) | (0.203-0.21) | (0.208-0.215) |
| **Carson 1991** | (0.033-0.034) | (0.065-0.067) | (0.021-0.022) | (0.019-0.02) | (0.093-0.097) | (0.115-0.119) | (0.117-0.121) | (0.121-0.125) |
| **Carson 1993** | (0.032-0.034) | (0.063-0.066) | (0.02-0.021) | (0.019-0.02) | (0.092-0.096) | (0.114-0.118) | (0.116-0.12) | (0.12-0.124) |
| **Carson 1995** | (0.032-0.033) | (0.062-0.064) | (0.021-0.022) | (0.019-0.02) | (0.091-0.094) | (0.112-0.116) | (0.117-0.121) | (0.121-0.125) |
| **Ebbetts 1995** |  | (0.044-0.046) | (0.029-0.03) | (0.026-0.028) | (0.078-0.081) | (0.1-0.103) | (0.139-0.144) | (0.143-0.148) |
| **Gaylor 1995** | 0.045 |  | (0.06-0.062) | (0.058-0.06) | (0.052-0.054) | (0.074-0.077) | (0.175-0.18) | (0.179-0.185) |
| **Da ek dow go et 1993** | 0.03 | 0.061 |  | (0.014-0.014) | (0.09-0.094) | (0.112-0.116) | (0.124-0.128) | (0.128-0.132) |
| **Da ek dow go et 1995** | 0.027 | 0.059 | 0.014 |  | (0.088-0.091) | (0.109-0.113) | (0.121-0.126) | (0.125-0.129) |
| **Mammoth 1993** | 0.079 | 0.053 | 0.092 | 0.089 |  | (0.036-0.038) | (0.209-0.215) | (0.213-0.219) |
| **Mammoth 1995** | 0.101 | 0.075 | 0.114 | 0.111 | 0.037 |  | (0.235-0.242) | (0.24-0.247) |
| **Rose 1995** | 0.141 | 0.177 | 0.126 | 0.123 | 0.212 | 0.239 |  | (0.013-0.013) |
| **Rose 1993** | 0.145 | 0.182 | 0.13 | 0.127 | 0.216 | 0.243 | 0.013 |  |
| **Moriah 1995** | 0.049 | 0.018 | 0.065 | 0.062 | 0.051 | 0.074 | 0.182 | 0.186 |
| **Reynolds 1993** | 0.022 | 0.054 | 0.031 | 0.029 | 0.087 | 0.109 | 0.143 | 0.146 |
| **Saddlebag 1991** | 0.032 | 0.039 | 0.033 | 0.03 | 0.071 | 0.092 | 0.138 | 0.142 |
| **Sonora 1995** | 0.03 | 0.047 | 0.049 | 0.046 | 0.084 | 0.107 | 0.166 | 0.17 |
| **Sonora 1991** | 0.02 | 0.038 | 0.038 | 0.035 | 0.074 | 0.096 | 0.154 | 0.159 |
| **Sweetwaters 1995** | 0.072 | 0.092 | 0.09 | 0.087 | 0.131 | 0.156 | 0.214 | 0.219 |
| **Sweetwaters 1993** | 0.033 | 0.051 | 0.052 | 0.049 | 0.089 | 0.111 | 0.171 | 0.175 |
| **Sweetwaters 1992** | 0.036 | 0.053 | 0.055 | 0.052 | 0.092 | 0.115 | 0.174 | 0.178 |
| **Tioga 1995** | 0.043 | 0.015 | 0.059 | 0.056 | 0.051 | 0.074 | 0.173 | 0.178 |

| Table S5. Pairwise G_ST_ between collection years and sites. Observed G_ST_ values are show below the diagonal while bootstrapped 95% confidence intervals are shown above the diagonal. Continued from Table S3. | | | | | | | | | |
| --- | --- | --- | --- | --- | --- | --- | --- | --- | --- |
|  | **Moriah 1995** | **Reynolds 1993** | **Saddlebag 1991** | **Sonora 1995** | **Sonora 1991** | **Sweetwaters 1995** | **Sweetwaters 1993** | **Sweetwaters 1992** | **Tioga 1995** |
| **Castle 1991** | (0.163-0.168) | (0.13-0.134) | (0.124-0.128) | (0.152-0.157) | (0.14-0.145) | (0.197-0.203) | (0.156-0.16) | (0.159-0.164) | (0.158-0.163) |
| **Castle 1992** | (0.195-0.201) | (0.161-0.166) | (0.154-0.159) | (0.183-0.189) | (0.171-0.176) | (0.229-0.236) | (0.186-0.192) | (0.19-0.196) | (0.19-0.196) |
| **Castle 1993** | (0.159-0.164) | (0.126-0.13) | (0.119-0.123) | (0.148-0.152) | (0.136-0.141) | (0.193-0.199) | (0.151-0.156) | (0.155-0.159) | (0.154-0.158) |
| **Castle 1994** | (0.187-0.192) | (0.152-0.157) | (0.146-0.151) | (0.175-0.18) | (0.163-0.168) | (0.22-0.226) | (0.178-0.183) | (0.181-0.187) | (0.181-0.187) |
| **Castle 1995** | (0.16-0.165) | (0.127-0.131) | (0.12-0.124) | (0.149-0.153) | (0.137-0.141) | (0.194-0.2) | (0.153-0.158) | (0.156-0.161) | (0.155-0.16) |
| **Carson 1991** | (0.068-0.07) | (0.034-0.035) | (0.029-0.03) | (0.053-0.055) | (0.044-0.045) | (0.095-0.098) | (0.057-0.059) | (0.06-0.063) | (0.062-0.065) |
| **Carson 1993** | (0.067-0.07) | (0.033-0.035) | (0.029-0.03) | (0.052-0.054) | (0.043-0.044) | (0.094-0.097) | (0.056-0.058) | (0.059-0.061) | (0.061-0.064) |
| **Carson 1995** | (0.065-0.067) | (0.033-0.034) | (0.029-0.03) | (0.052-0.053) | (0.042-0.043) | (0.093-0.096) | (0.056-0.058) | (0.059-0.061) | (0.059-0.062) |
| **Ebbetts 1995** | (0.048-0.049) | (0.022-0.022) | (0.031-0.032) | (0.03-0.031) | (0.02-0.02) | (0.071-0.074) | (0.032-0.034) | (0.036-0.037) | (0.042-0.044) |
| **Gaylor 1995** | (0.018-0.018) | (0.053-0.055) | (0.038-0.039) | (0.046-0.048) | (0.037-0.039) | (0.09-0.093) | (0.05-0.052) | (0.052-0.055) | (0.015-0.015) |
| **Da ek dow go et 1993** | (0.064-0.066) | (0.03-0.032) | (0.032-0.033) | (0.048-0.05) | (0.038-0.039) | (0.089-0.092) | (0.051-0.053) | (0.054-0.056) | (0.058-0.06) |
| **Da ek dow go et 1995** | (0.06-0.063) | (0.028-0.029) | (0.029-0.031) | (0.045-0.047) | (0.035-0.036) | (0.086-0.089) | (0.048-0.05) | (0.051-0.053) | (0.055-0.057) |
| **Mammoth 1993** | (0.05-0.052) | (0.085-0.089) | (0.069-0.072) | (0.083-0.086) | (0.073-0.075) | (0.129-0.134) | (0.087-0.091) | (0.091-0.094) | (0.05-0.052) |
| **Mammoth 1995** | (0.072-0.075) | (0.107-0.111) | (0.091-0.094) | (0.105-0.109) | (0.094-0.098) | (0.153-0.158) | (0.109-0.113) | (0.113-0.117) | (0.072-0.075) |
| **Rose 1995** | (0.179-0.185) | (0.14-0.145) | (0.136-0.14) | (0.164-0.169) | (0.152-0.157) | (0.211-0.217) | (0.168-0.173) | (0.171-0.177) | (0.171-0.176) |
| **Rose 1993** | (0.183-0.189) | (0.144-0.149) | (0.14-0.145) | (0.168-0.173) | (0.156-0.161) | (0.215-0.222) | (0.172-0.177) | (0.176-0.181) | (0.175-0.181) |
| **Moriah 1995** |  | (0.056-0.059) | (0.04-0.042) | (0.05-0.052) | (0.041-0.042) | (0.094-0.098) | (0.054-0.056) | (0.057-0.059) | (0.017-0.018) |
| **Reynolds 1993** | 0.057 |  | (0.035-0.036) | (0.038-0.039) | (0.028-0.029) | (0.079-0.082) | (0.041-0.042) | (0.044-0.045) | (0.051-0.052) |
| **Saddlebag 1991** | 0.041 | 0.036 |  | (0.043-0.045) | (0.034-0.035) | (0.085-0.089) | (0.047-0.049) | (0.05-0.052) | (0.035-0.037) |
| **Sonora 1995** | 0.051 | 0.038 | 0.044 |  | (0.02-0.021) | (0.075-0.078) | (0.036-0.037) | (0.039-0.04) | (0.044-0.046) |
| **Sonora 1991** | 0.041 | 0.029 | 0.034 | 0.021 |  | (0.064-0.067) | (0.025-0.026) | (0.028-0.029) | (0.035-0.036) |
| **Sweetwaters 1995** | 0.096 | 0.081 | 0.087 | 0.076 | 0.065 |  | (0.048-0.049) | (0.051-0.053) | (0.088-0.091) |
| **Sweetwaters 1993** | 0.055 | 0.042 | 0.048 | 0.036 | 0.025 | 0.048 |  | (0.011-0.011) | (0.048-0.05) |
| **Sweetwaters 1992** | 0.058 | 0.045 | 0.051 | 0.04 | 0.029 | 0.052 | 0.011 |  | (0.051-0.053) |
| **Tioga 1995** | 0.017 | 0.051 | 0.036 | 0.045 | 0.036 | 0.09 | 0.049 | 0.052 |  |

| Table S6. Summary statistics for linear model predicting geographic distance using G_ST_. | | | |
| --- | --- | --- | --- |
|  | **Estimate** | **Standard error** | **t-value** |
| Intercept | 24.42 | 2.79 | 8.74 |
| G_ST_ | 565.73 | 18.77 | 30.15 |
